# Light ‘em up: efficient screening of gold grids in cryo-EM

**DOI:** 10.1101/2022.04.27.489675

**Authors:** Wim J.H. Hagen

## Abstract

Transmission electron cryo-microscopy (cryo-EM) allows for obtaining 3D structural information by imaging macromolecules embedded in thin layers of amorphous ice. To obtain high-resolution structural information, samples need to be thin to minimize inelastic scattering which blurs images. During data collection sessions, time spent on finding areas on the cryo-EM grid with optimal ice thickness should be minimized as imaging time on high-end Transmission Electron Microscope TEM systems is costly. Recently, grids covered with thin gold films have become popular due to their stability and reduced beam-induced motion of the sample. Gold foil grids have substantially different densities between the gold foil and ice, effectively resulting in the loss of dynamic range between thin and thick regions of ice, making it challenging to find areas with suitable ice thickness efficiently during grid screening and thus increase expensive imaging time. Here, an energy filter-based plasmon imaging is presented as a fast and easy method for grid screening of the gold grids.

## Introduction

Structure determination by cryo-EM (Frank, 2006) ideally requires vitrified specimens with a thin frozen-hydrated layer only slightly thicker than the structure of interest, since final image resolution has a direct correlation with ice thickness (Rheinberger et al., 2021; Rice et al., 2018). The sample carrier typically used for vitrification of cryo-EM grids is a metal grid covered with a thin electron-transparent carbon film containing a pattern of holes “holey-carbon grid” (Ermantraut et al., 1998; Quispe et al., 2007) that are to be filled with thin ice during the vitrification process. For this, a thin layer of sample is applied to the grid and then flash-frozen into a cryogen (Adrian et al., 1984). Although the vitrification process has seen several improvements in recent years (Dandey et al., 2018; Jain et al., 2012; Kontziampasis et al., 2019; Rubinstein et al., 2019), controlling ice thickness remains challenging and screening of grids in the TEM is still required. Screening involves acquiring image montages at low magnification to visualize the entire grid area, optionally followed by images at a medium magnification at selected positions expected to have optimal ice thickness (Carragher et al., 2000; Suloway et al., 2005); particle concentration is subsequently inspected in those selected areas. Several software packages for automated data acquisition, both open source and commercial, exist to aid this screening process (Tan et al., 2016) and the usage strategies of these software packages are similar. Screening typically starts with acquiring a full overview of the EM grid at very low magnification, followed by images or montages of single grid squares at medium magnification, such that holes in the film with sizes of several micrometers can be distinguished. On any TEM system, this requires using 1) the lowest possible magnification in LM mode (the objective lens is switched off) for mapping the entire grid, and 2) using a medium magnification LM mode for imaging or montaging a single grid square or using a low magnification Selected Area SA mode (objective lens switched on) to visualise grid squares. A microscope user will then visually inspect the images to find areas with thin ice based on how dark or light the holes look. Similarly, software packages for automatic data acquisition use grey values of the holes in an image to allow a user to set a filter to select holes within a certain ice-thickness range. This procedure has been very successful for samples prepared on standard holey carbon grids and greatly speeds up the acquisition session setup.

One of many limitations of cryo-EM structure determination has been beam-induced particle movement, in which particles in thin ice move around during exposure to an electron beam. This limitation has been minimized by the development of the so-called “gold grids” (Naydenova et al., 2020; Russo and Passmore, 2016, 2014). Gold grids use a gold carrier mesh grid covered with a thin gold film with patterned holes. Compared to the historically used holey carbon films, thin gold films are considerably less electron transparent, while thin ice in the holes of the film is similarly electron transparent. This leads to a larger contrast range in images of gold grids. The larger contrast difference in the TEM requires some image processing filter or processing to judge relative ice thickness in the holes. The design of such a filtering scheme should be based on how electrons differently interact with the sample.

When electrons used for imaging pass through the sample, several interactions take place with the sample: no scattering, elastic scattering, and inelastic scattering. Electrons that are inelastically scattered lose energy and will be focussed differently by the downstream optics of the microscope leading to image blurring. Inelastic scattering depends on sample thickness: the thicker the sample, the larger the chance that electrons will be scattered once, or multiple times while passing through the sample. The resulting blurring effect can be reduced by using an energy filter, which is a dispersive prism that separates electrons by the energy they possess, followed by an energy selecting slit to only let through elastically scattered electrons that have not lost energy passing through the sample. This is called Zero Loss Imaging. The energy selecting slit can also be offset to only let certain energy loss ranges pass, e.g. the plasmon regime. Plasmons are inelastic scattering events with low energy loss (< 50 eV). In the plasmon regime, vitreous, cubic, and hexagonal ice show a broad plasmon peak around 23 eV (Leapman and Sun, 1995), while gold shows narrow plasmon peaks at very low energy losses, buried inside the tail of the zero-loss peak as seen on non-monochromated TEM systems (Schaffer et al., 2010). Positioning an energy slit around 20 eV energy loss, with an energy slit width of 20 eV to block the zero-loss electrons, produces images with a smaller image contrast range, which simplifies the visual interpretation of ice thickness in the holes.

Additionally, using an objective aperture limits the energy filter’s collection angle, further decreasing the contrast range of the plasmon images. Under low electron exposure conditions, the intensity of plasmon images is low enough to use counting mode when using direct detectors, which minimizes background noise in the acquired images, while increased specimen thickness will lead to more plasmon events. A properly aligned microscope and a properly tuned energy filter allow identification of relative ice thickness at even the lowest microscope magnifications with very low exposure dose. The nature of the plasmon mechanism does not allow for distinguishing between vitreous, cubic, or hexagonal ice, it only indicates relative thickness, albeit with great detail. Several methods have been published to assist microscope operators with ice thickness determination, mainly based on other variations of energy-filter usage (Rheinberger et al., 2021; Rice et al., 2018). Plasmon imaging provides an additional method especially suitable for screening gold grids on TEM systems equipped with an energy filter.

## Methods

Assuming a properly aligned microscope and properly tuned energy filter, the only setting needed is a 15 eV energy slit width and 20 eV energy loss offset. On Thermo Scientific Selectris(X) energy filters, the energy loss offset can be achieved either using the magnetic prism (energy shift) or by the energy filter control panel changing the high tension (HT offset). For certain types of Gatan energy filters, HT offset is not available and the two options then are energy shift or using the drift tube. All options can be done in most currently available software packages as shown in Figure 1.

**Figure 1.**
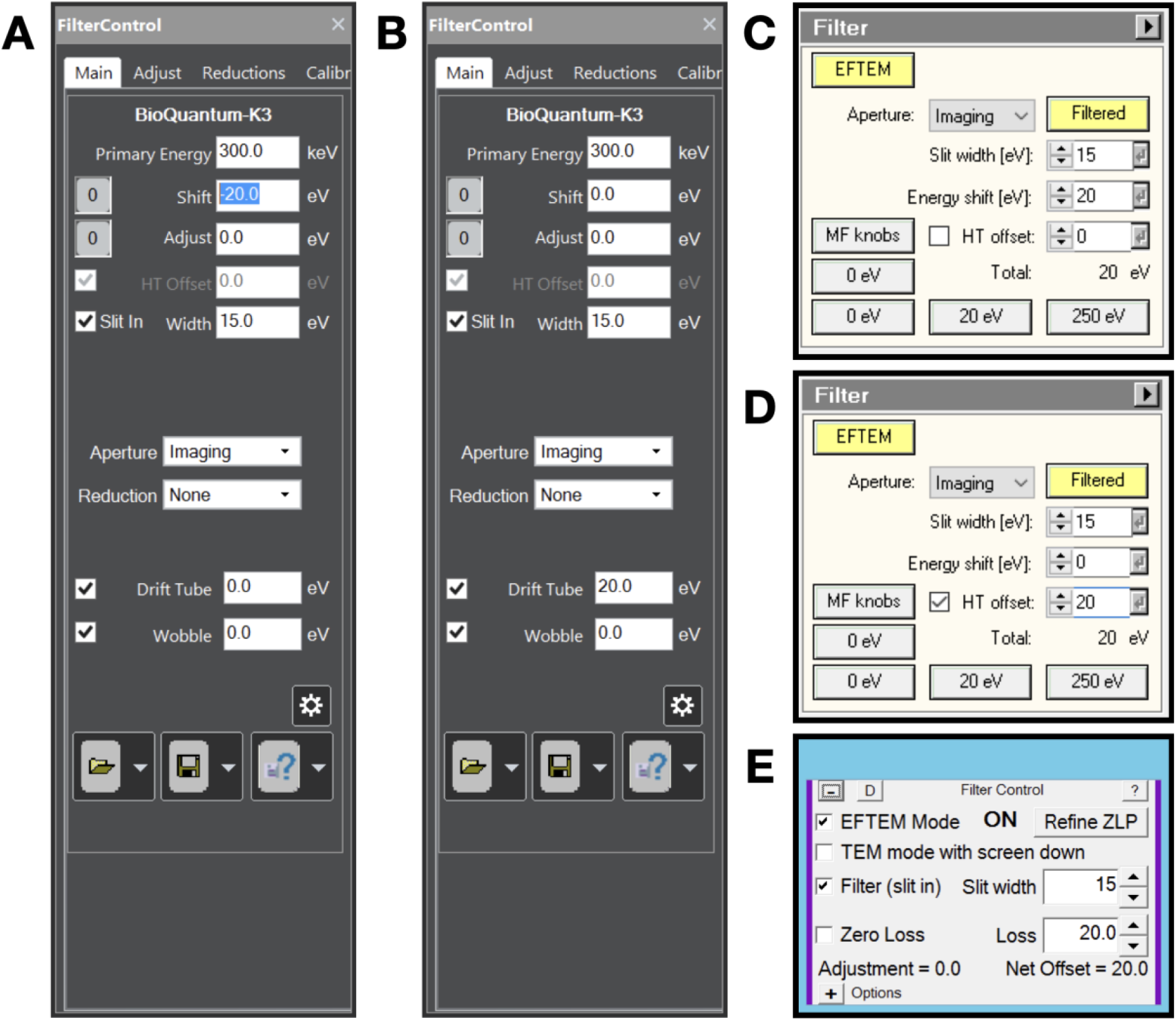
Energy filter settings for plasmon imaging when A) using a Gatan energy filter controlled by Digital Micrograph software and using energy shift, B) using a Gatan energy filter controlled by Digital Micrograph software and using Drift tube offset, C) using a Thermo Scientific Selectris(X) energy filter using energy shift, D) using a Thermo Scientific Selectris(X) energy filter using high tension offset and E) using any energy filter through SerialEM.

In typical cryo-TEM systems, optimal alignment of the microscope, especially for the lower magnification ranges is a condition rarely met. Several alignments and energy filter settings need to be tweaked, which might not always be possible for the user as it might require service credentials from the microscope vendor. One can, however, work around these issues through the many available data acquisition software packages. All steps required are described next. The minimum starting point is that EFTEM LM and EFTEM SA mode are properly aligned by the microscope vendor, meaning that the energy filter can image at all low magnifications without the differential pumping aperture between the column and the projection chamber blocking the field of view. Using the plasmon regime requires an energy filter to be properly tuned at several different magnifications. The currently most popular energy filters are made by Gatan, whose Digital Micrograph software has a GIF Settings Manager to store energy filter tuning per microscope magnification. This feature needs to be activated and set up by the Gatan service engineer upon installation of the energy filter.

Thermo Scientific Selectris(X) energy filters store tuning values per magnification by default. The energy filter needs to be tuned for each magnification intended to be used for plasmon imaging. Any energy loss imaging has to be done with the sample in focus as defocus leads to image blur. This is more important for SA mode (objective lens on), where tens of micro-meter focus offset lead to extreme blurring, than for LM mode (objective lens off), where hundreds of micro-meter focus offset lead to extreme blurring. Setting correct focus and fixing image astigmatism in LM mode is often overlooked during alignment of the microscope, but it is easy to fix using available data acquisition software:

- Load a vitrified gold grid into the microscope.
- Set the stage to eucentric height (minimum image movement when wobbling specimen tilt using the goniometer).
- Set the energy filter to 15 eV energy slit width and 20 eV energy loss.
- Using live imaging in any capable camera software, adjust defocus and image astigmatism until the image appears sharp. On Thermo Scientific TEM systems, in LM mode the diffraction stigmator acts as the image stigmator.

On Thermo Scientific TEM systems, the stigmator settings are automatically saved as user settings, and the defocus setting needs to be written down to be used as an offset to be applied in the data acquisition software used. When using LM mode magnification, one can additionally use an SA aperture (in LM mode, the SA aperture acts as the objective aperture). When using a low SA mode magnification, one can use a standard objective aperture. An aperture increases the visible contrast difference of varying ice thickness.

## Results

Most modern cryo-EM systems fitted with energy filters have direct detectors. Such detectors typically have two operation modes, linear (integration) mode for high exposure dose rate imaging, and (electron) counting mode for low exposure dose rate imaging. When imaging using a direct detector without an energy selecting slit, or in zero-loss mode, the high dose-rate reaching the detector requires the use of linear mode, or a lower dose rate in counting mode, requiring long exposure times to achieve acceptable imaging conditions. Plasmon imaging shows empty holes as black since no sample is present and interaction of electrons with the sample is completely absent. Holes covered with thin ice show as signal, with signal level increasing with sample thickness, even in images acquired at the very lowest magnification that can be used on the microscope. Since the plasmon signal is very low, counting mode can be used, which minimizes the background noise of the images. One can additionally use an aperture to improve imaging contrast, which further increases the visibility of ice thickness differences. The differences between the three strategies (LM grid mapping, LM square-mapping, SA square-mapping) typically used when inspecting grids are shown for grid LM mapping (Figure 2), visualizing a grid square using a single image (Figure 3), and using image montages using a low SA magnification (Figure 4).

**Figure 2.**
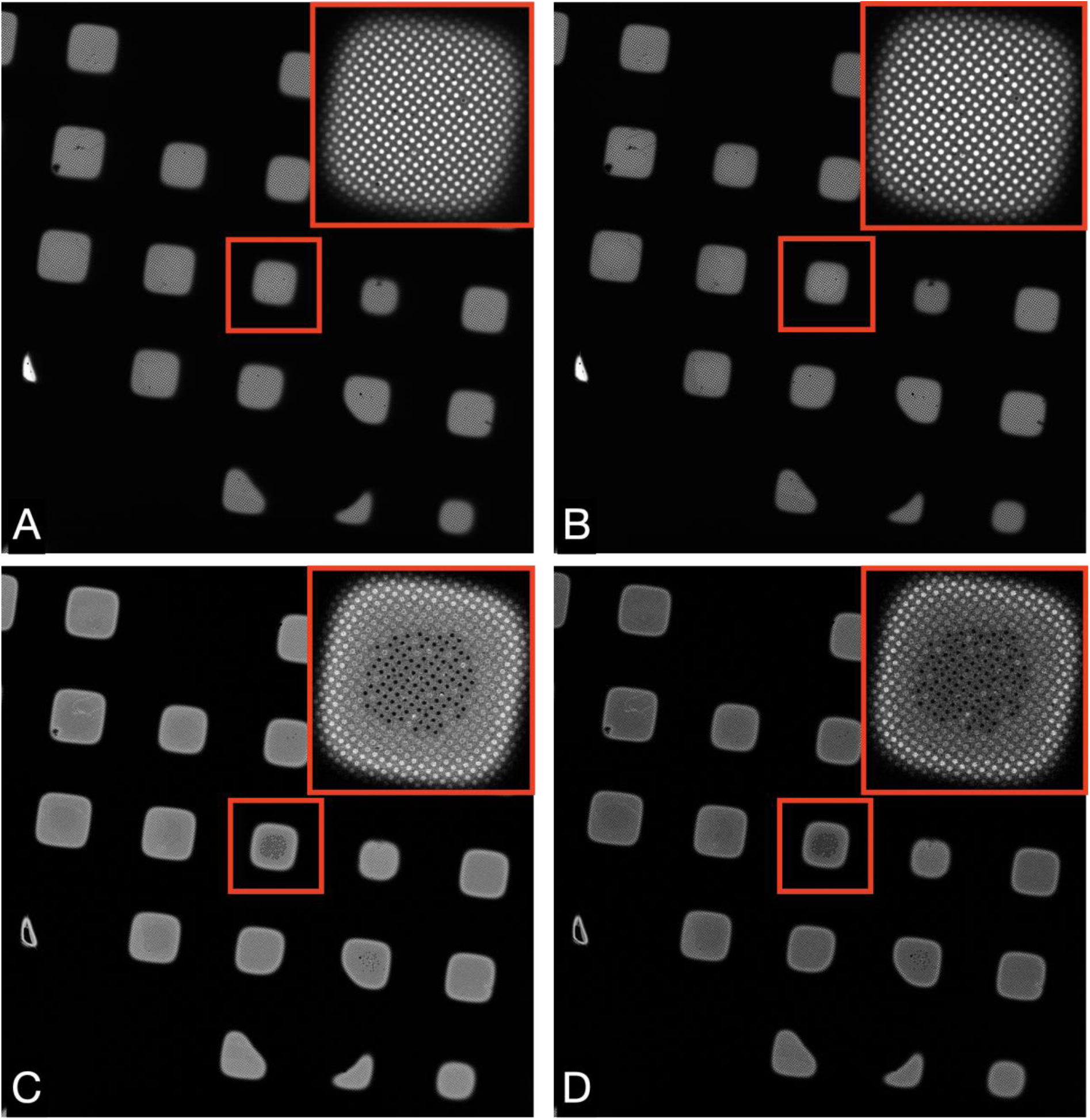
Images of a Quantifoil UltrAuFoil 1.2/1.3 grid, acquired on a Thermo Scientific Titan Krios G1with Gatan Bioquantum K2 detector using EFTEM LM mode 135x magnification (field of view 418 × 433 μm), 1-second exposure, total dose 0.001 e/Å^2^. The upper right corners show a magnified inset of the marked central square. Image contrast/brightness has not been changed manually. A) Unfiltered image, B) Zero loss filtered linear mode image, C) plasmon counting mode image, and D) plasmon counting mode image with 40 μm SA aperture inserted.

**Figure 3.**
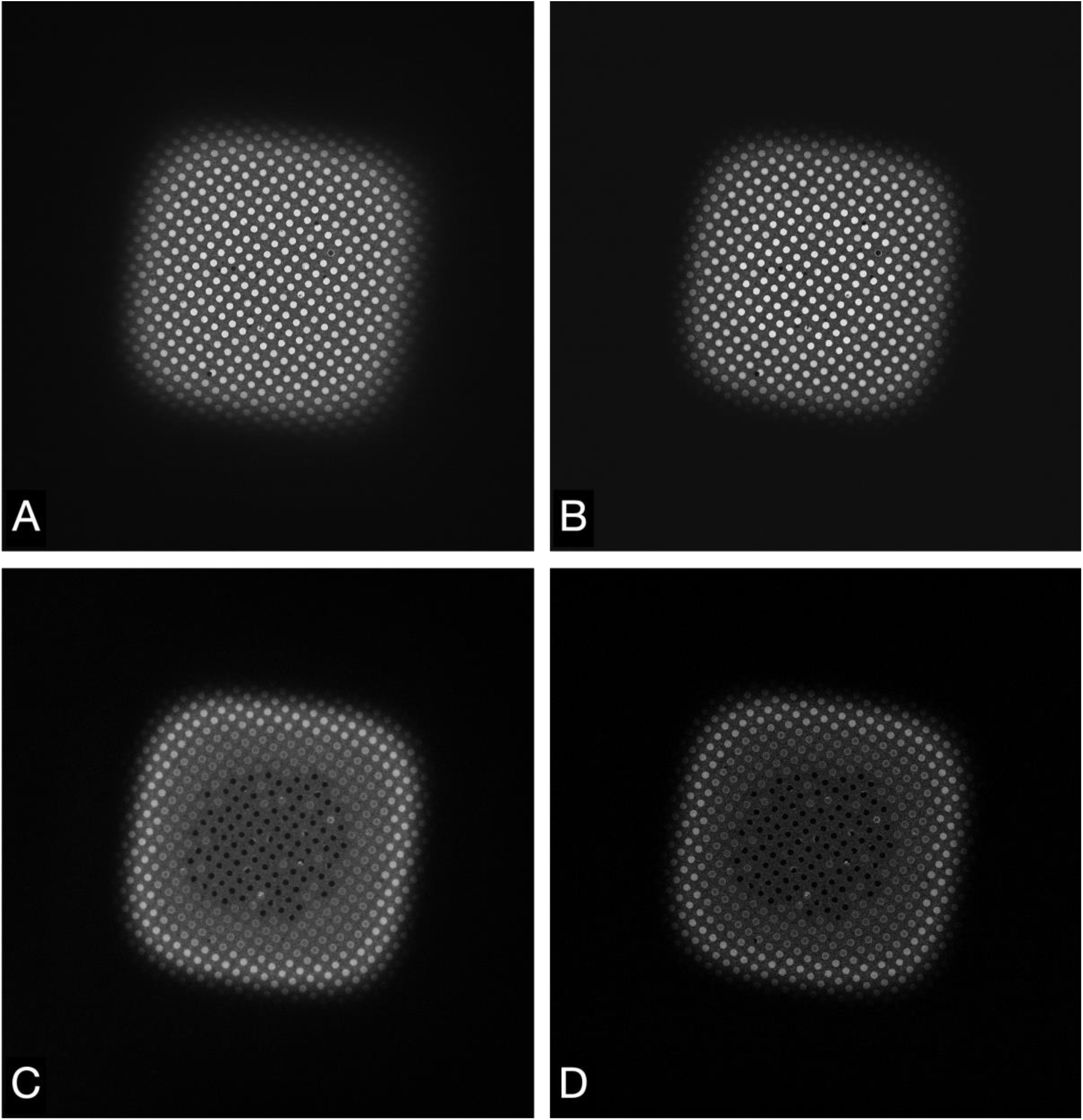
Images of a Quantifoil UltrAuFoil 1.2/1.3 grid square, acquired on a Thermo Scientific Titan Krios G1with Gatan Bioquantum K2 detector using EFTEM LM mode 740x magnification (field of view 67 × 69 μm), 1-second exposure, total dose 0.003 e/Å^2^. Image contrast/brightness has not been changed manually. A) Unfiltered image, B) Zero loss filtered linear mode image, C) plasmon counting mode image, and D) plasmon counting mode image with 40 μm SA aperture inserted (lower right).

**Figure 4.**
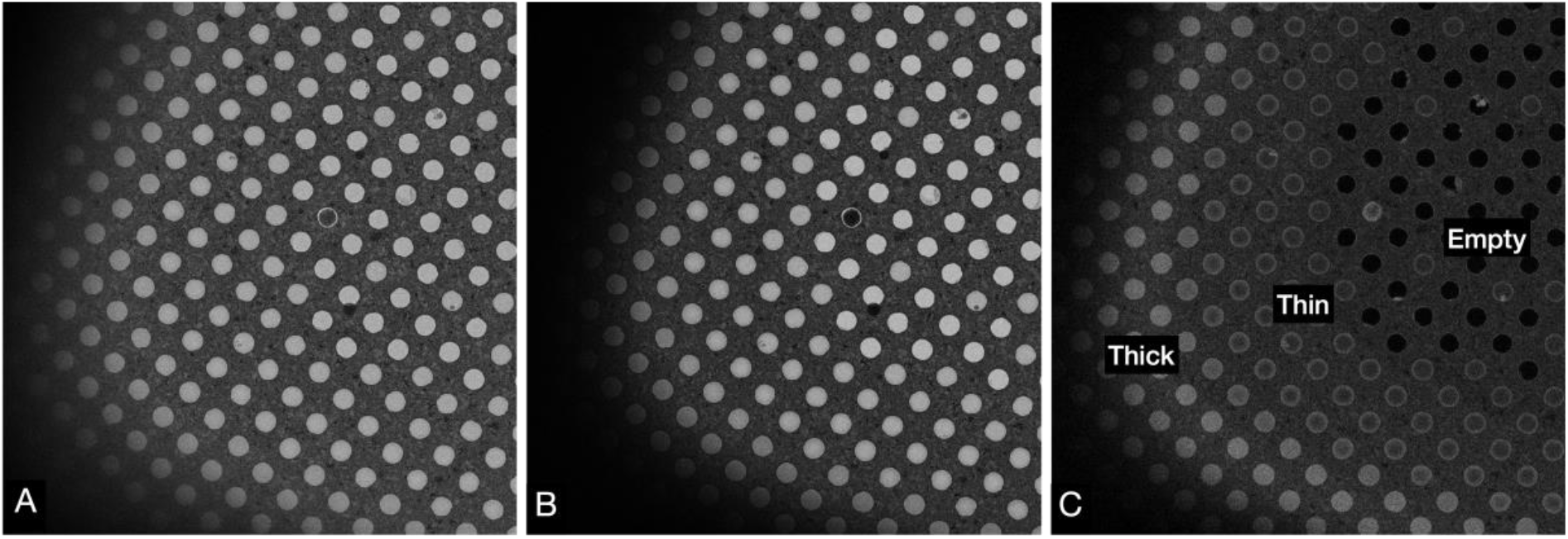
Images of a Quantifoil UltrAuFoil 1.2/1.3 grid, acquired on a Thermo Scientific Titan Krios G1with Gatan Bioquantum K2 detector using EFTEM SA mode 2250x magnification (field of view 23 × 24 μm) with a 70 μm objective aperture inserted, 1-second exposure, total dose 0.003 e/Å^2^. Image contrast/brightness has not been changed manually. A) Unfiltered image with -100 μm defocus, B) Zero loss filtered linear mode image with -100 μm defocus, and C) plasmon counting mode image without any defocus(right).

## Discussion

A method has been described to assist microscope users in quickly judging relative ice thickness for cryo-EM data acquisition. It requires an energy filter but no additional images and processing steps that are typically required to produce thickness maps (Rheinberger et al., 2021). However, the described method only provides relative thickness information whereas thickness maps provide absolute thickness, albeit with less detail about any ice gradients within single holes in the gold film. I hope this simple and readily available plasmon method helps cryo-EM users in choosing optimal areas for imaging on vitrified gold grids. Additionally, this method could open new possibilities by skipping the grid square step altogether by targeting holes directly from the lowest magnification montages.

## Acknowledgments

I thank Sonja Welsch, Svetlana Dodonova and William Wan for their comments.

## References

Adrian, M., Dubochet, J., Lepault, J., McDowall, A.W., 1984. Cryo-electron microscopy of viruses. Nature 308, 32–36. https://doi.org/10.1038/308032a0

Carragher, B., Kisseberth, N., Kriegman, D., Milligan, R.A., Potter, C.S., Pulokas, J., Reilein, A., 2000. Leginon: an automated system for acquisition of images from vitreous ice specimens. J. Struct. Biol. 132, 33–45. https://doi.org/10.1006/jsbi.2000.4314

Dandey, V.P., Wei, H., Zhang, Z., Tan, Y.Z., Acharya, P., Eng, E.T., Rice, W.J., Kahn, P.A., Potter, C.S., Carragher, B., 2018. Spotiton: New features and applications. J. Struct. Biol. 202, 161–169. https://doi.org/https://doi.org/10.1016/j.jsb.2018.01.002

Ermantraut, E., Wohlfart, K., Tichelaar, W., 1998. Perforated support foils with pre-defined hole size, shape and arrangement. Ultramicroscopy 74, 75–81. https://doi.org/10.1016/S0304-3991(98)00025-4

Frank, J., 2006. Three-Dimensional Electron Microscopy of Macromolecular Assemblies: Visualization of Biological Molecules in Their Native State. Oxford University Press, New York. https://doi.org/10.1093/acprof:oso/9780195182187.001.0001

Jain, T., Sheehan, P., Crum, J., Carragher, B., Potter, C.S., 2012. Spotiton: A prototype for an integrated inkjet dispense and vitrification system for cryo-TEM. J. Struct. Biol. 179, 68–75. https://doi.org/https://doi.org/10.1016/j.jsb.2012.04.020

Kontziampasis, D., Klebl, D.P., Iadanza, M.G., Scarff, C.A., Kopf, F., Sobott, F., Monteiro, D.C.F., Trebbin, M., Muench, S.P., White, H.D., 2019. A cryo-EM grid preparation device for time-resolved structural studies. IUCrJ 6, 1024–1031. https://doi.org/10.1107/S2052252519011345

Leapman, R.D., Sun, S., 1995. Cryo-electron energy loss spectroscopy: observations on vitrified hydrated specimens and radiation damage. Ultramicroscopy. https://doi.org/10.1016/0304-3991(95)00019-W

Naydenova, K., Jia, P., Russo, C.J., 2020. Cryo-EM with sub–1 Å specimen movement. Science (80-.). 370, 223–226. https://doi.org/10.1126/SCIENCE.ABB7927

Quispe, J., Damiano, J., Mick, S.E., Nackashi, D.P., Fellmann, D., Ajero, T.G., Carragher, B., Potter, C.S., 2007. An improved holey carbon film for cryo-electron microscopy. Microsc. Microanal. 13, 365–371. https://doi.org/10.1017/S1431927607070791

Rheinberger, J., Oostergetel, G., Resch, G.P., Paulino, C., 2021. Optimized cryo-EM data-acquisition workflow by sample-thickness determination. Acta Crystallogr. Sect. D, Struct. Biol. 77, 565–571. https://doi.org/10.1107/S205979832100334X

Rice, W.J., Cheng, A., Noble, A.J., Eng, E.T., Kim, L.Y., Carragher, B., Potter, C.S., 2018. Routine determination of ice thickness for cryo-EM grids. J. Struct. Biol. 204, 38–44. https://doi.org/10.1016/j.jsb.2018.06.007

Rubinstein, J.L., Guo, H., Ripstein, Z.A., Haydaroglu, A., Au, A., Yip, C.M., Di Trani, J.M., Benlekbir, S., Kwok, T., 2019. Shake-it-off: a simple ultrasonic cryo-EM specimen-preparation device. Acta Crystallogr. Sect. D 75, 1063–1070. https://doi.org/10.1107/S2059798319014372

Russo, C.J., Passmore, L.A., 2016. Ultrastable gold substrates: Properties of a support for high-resolution electron cryomicroscopy of biological specimens. J. Struct. Biol. 193, 33–44. https://doi.org/10.1016/J.JSB.2015.11.006

Russo, C.J., Passmore, L.A., 2014. Ultrastable gold substrates for electron cryomicroscopy. Science (80-.). 346, 1377–1380. https://doi.org/10.1126/SCIENCE.1259530

Schaffer, B., Grogger, W., Kothleitner, G., Hofer, F., 2010. Comparison of EFTEM and STEM EELS plasmon imaging of gold nanoparticles in a monochromated TEM. Ultramicroscopy 110, 1087–1093. https://doi.org/https://doi.org/10.1016/j.ultramic.2009.12.012

Suloway, C., Pulokas, J., Fellmann, D., Cheng, A., Guerra, F., Quispe, J., Stagg, S., Potter, C.S., Carragher, B., 2005. Automated molecular microscopy: the new Leginon system. J. Struct. Biol. 151, 41–60. https://doi.org/10.1016/j.jsb.2005.03.010

Tan, Y.Z., Cheng, A., Potter, C.S., Carragher, B., 2016. Automated data collection in single particle electron microscopy. Reprod. Syst. Sex. Disord. 65, 43–56. https://doi.org/10.1093/jmicro/dfv369

